# The p300 KAT inhibitor, A-485, inhibits melanogenesis and melanin content in human melanocytes

**DOI:** 10.1101/2024.02.14.580089

**Authors:** Nicole Grbic, Aaron Richard Waddell, Kassidy Leibowitz, W. Austin Wyant, Marianne Collard, Rhoda M. Alani

## Abstract

Hyperpigmentation disorders are commonly diagnosed dermatologic conditions that can be cosmetically distressing for patients and cause negative psychosocial impacts. There is a need to better understand the underlying pathophysiology of pigmentary disorders as well as develop improvements in the management of these disorders. Here, we evaluated a p300 (CBP/p300) histone acetyltransferase (HAT) inhibitor, A-485, to determine if epigenetically targeting melanogenesis could be of therapeutic value. We find that A-485 treatment of primary human melanocytes and SK-MEL-30 melanoma cells drastically reduces the mRNA and protein expression of MITF and DCT, genes involved in melanin synthesis, and that this is accompanied by a reduction in melanin content. These results suggest that epigenetically targeting melanin synthesis with the A-485 p300 HAT inhibitor may be beneficial for the treatment of hyperpigmentation disorders.

## Introduction

Pigmentary disorders commonly present in patients and are of concern beyond physical appearance due to the negative psychosocial impacts reported by patients (Taylor et al., 2008). Pigmentary disorders are one of the top five most common dermatologic conditions diagnosed in African Americans (Alexis et al., 2007) and include post-inflammatory hyperpigmentation, melasma, solar lentigines, ephelides, and café au lait macules (Plensdorf & Martinez, 2009). The US Census Bureau has predicted that by 2060, more than half of the US population will have skin of color (Bureau, 2012); it is therefore critical for the scientific community to better understand the underlying pathophysiology and management of pigmentary disorders.

Hyperpigmentation disorders are often due to increased melanogenesis, which includes the production of melanin in the melanosome and the transport of melanin from melanocytes to keratinocytes (Pavel, 1993). Although there is no racial difference in the number of melanocytes present (Staricco & Pinkus, 1957), darkly pigmented skin contains greater total melanin content (Smit et al., 1998) due to an increased number of melanosomes that are larger in size and more individually dispersed compared to lightly pigmented skin (Szabo et al., 1969). The increased amount of melanin in more pigmented skin has been proposed to increase the risk of pigmentary disorders, including post-inflammatory hyperpigmentation (Vashi & Kundu, 2013).

Microphthalmia-associated transcription factor (MITF) is an essential transcription factor for melanin production which also regulates melanocyte development, survival, and proliferation (Vachtenheim & Borovansky, 2010). MITF promotes transcription of melanogenesis enzymes including tyrosinase (McIntyre et al.), the rate-limiting enzyme for melanin synthesis, dopachrome tautomerase (DCT, also known as tyrosine-related protein-2, TRP-2), and tyrosine-related protein 1 (TRP-1) (Vachtenheim & Borovansky, 2010). Moreover, MITF itself is critically regulated through epigenetic changes including DNA methylation and histone acetylation (Zhou et al., 2021). The transcriptional co-activators CREB-binding protein and p300 (CBP/p300) possess histone acetyltransferase (HAT) activity and play an important role in activating MITF and its downstream melanogenesis-related target genes (Zhou et al., 2021). Phosphorylated CREB recruits CBP/p300 to promote MITF transcription, while MITF recruits CBP/p300 to activate the transcription of key pigmentation genes including TYR, DCT, and TRP-1 (Zhou et al., 2021).

We have previously shown that inhibition of CBP/p300 activity with the potent and selective HAT inhibitor, A-485, suppresses growth in melanomas in an MITF-dependent fashion leading to inhibited expression of MITF and its downstream target, TRP-1 (Kim et al., 2019). We therefore hypothesized that A-485 would also inhibit cellular melanin content, which could be of potential clinical benefit in the setting of cutaneous hyperpigmentation.

## Results

To explore p300 HAT effects on cellular pigment content, SK-MEL-30 melanoma cells were incubated with 5 µM A-485 for 24 hours and MITF and DCT mRNA and protein levels were evaluated. Treatment of SK-MEL-30 melanoma cells with 5 µM A-485 led to reduced MITF and DCT mRNA (Figure 1A) and protein levels (Figure 1B) following 24 hours of treatment as well as significantly reduced cellular pigmentation following 6 days of treatment (Figure 1C, 1D) (p<0.005). Evaluation of A-485 effects on pigmentation in human primary melanocytes also led to reduced MITF and DCT mRNA (Figure 1E) and protein expression (Figure 1F) along with decreased melanin content following 6-days of treatment with A-485 (Figure 1G) (p<0.05).

**Figure 1:**
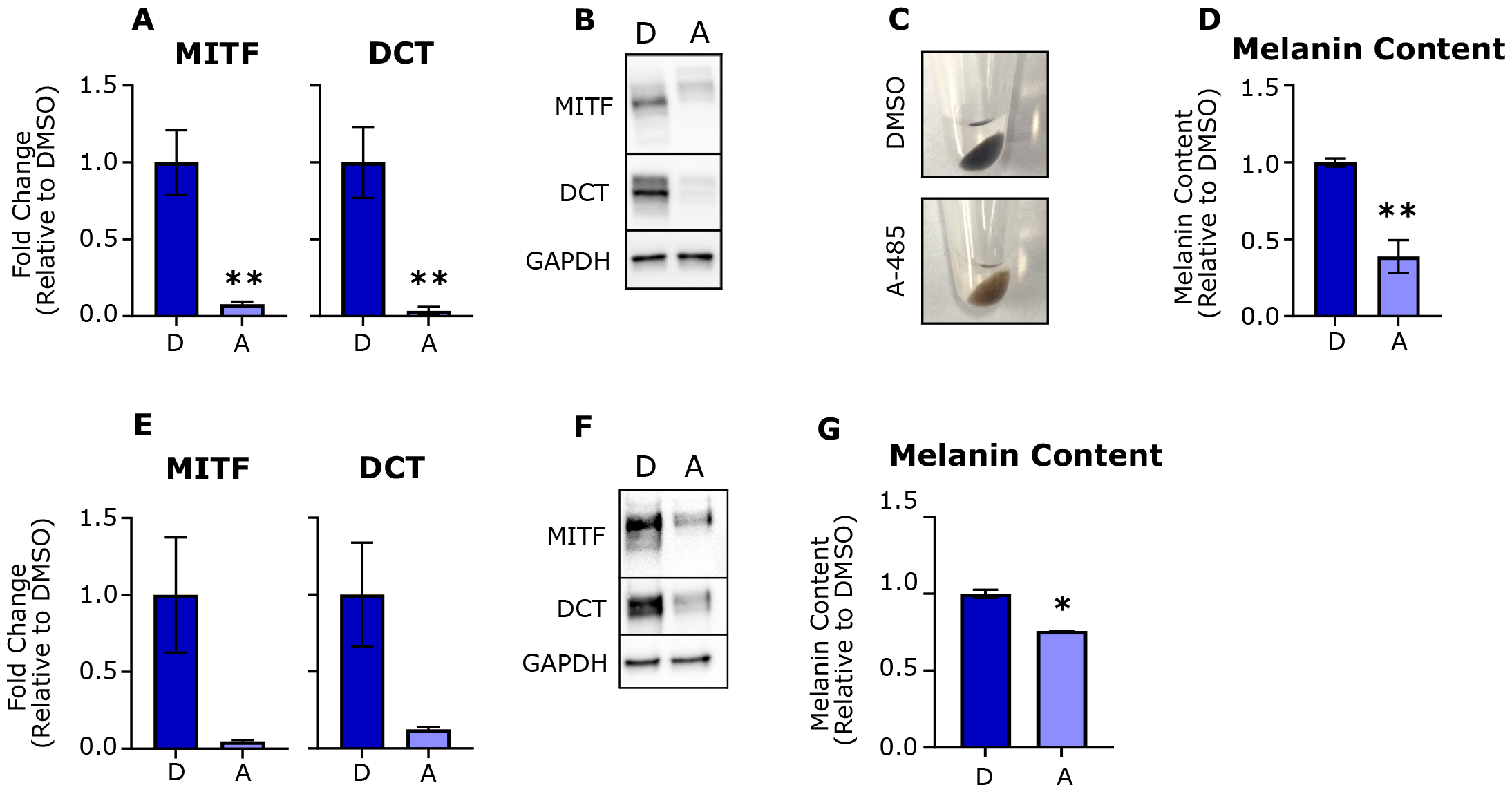
p300 KAT inhibitor A-485 attenuates expression of genes critical for melanogenesis, resulting in decreased pigmentation. (A) Expression of MITF and DCT were assayed via qPCR in SK-MEL-30 cells treated with 5 µM A-485 for 24 hours (n=3, error bars are mean +/-SD). (B) Protein levels of MITF and DCT were assayed via immunoblotting in SK-MEL-30 cells treated with 5 µM A-485 for 24 hours. (C) SK-MEL-30 cells were treated with DMSO (control) or 5 µM A-485 for 6-days. The cells were pelleted and imaged for melanin content. (D) SK-MEL-30 cells were treated with DMSO (control) or 5 µM A-485 for 6-days. Cells were lysed and melanin content was quantified by measuring absorbance at 405 nm and comparing measured values to a standard curve (n=3, error bars are mean +/-SD). (E) Expression of MITF and DCT were assayed via qPCR in human primary melanocytes treated with 5 µM A-485 for 24 hours (representative data for n=2, error bars are mean +/-SD). (F) Protein levels of MITF and DCT were assayed via immunoblotting in human primary melanocytes treated with 5 µM A-485 for 24 hours. (G) Human primary melanocytes were treated with DMSO (control) or 5 µM A-485 for 6-days. Cells were lysed and melanin content was quantified by measuring absorbance at 405 nm and comparing measured values to a standard curve (n=2, error bars are mean +/-SD). ^*^ p < 0.05, ^**^ p < 0.005. P-value was determined by Student’s t-test.

## Conclusion

These results confirm the important role for p300 HAT activity in the regulation of pigmentation-associated genes and support the potential use of p300 HAT inhibitors as therapeutic agents for the treatment of disorders of hyperpigmentation. Of note, epigenetic-targeting drugs such as A-485 have the capability of influencing entire transcriptional programs to reduce melanin synthesis which may revert cells to a low pigmentation phenotype. Our study supports that small molecules which target the epigenome should be further evaluated to treat hyperpigmentation disorders.

## Materials and Methods

### Cell culture

SK-MEL-30 melanoma cells were obtained from Dr. Anurag Singh (Boston University, Boston, MA, USA). Primary human melanocytes (PCS-200-013) were obtained from the American Type Culture Collection (ATCC) (Manassas, VA). Cells were grown in a humidified incubator at 37 °C with 5% CO2. Sk-MEL-30 cells were cultured in Dulbecco’s Modified Eagle Medium (Gibco, ThermoFisher Scientific, Grand Island, NY, USA) supplemented with 10% bovine calf serum (Gibco, ThermoFisher Scientific, Grand Island, NY, USA), 1% penicillin/streptomycin (Gibco, ThermoFisher Scientific, Grand Island, NY, USA), and 2mM L-glutamine (Gibco, ThermoFisher Scientific, Grand Island, NY, USA). Melanocytes were grown in Melanocyte Growth Media, which was comprised of Dermal Cell Basal Medium (PCS-200-030) supplemented with the Melanocyte Growth Kit (PCS-200-042) (ATCC, Manassas, VA).

### Compounds

A-485 was purchased from MedChemExpress (Monmouth Junction, NJ, USA) and diluted in DMSO to obtain a 10 mM stock solution.

### Immunoblotting

SK-MEL-30 or human primary melanocytes were treated with DMSO or A-485 (5 µM) for 24 hours. Whole-cell protein lysates were prepared in M-PER buffer (Thermo Scientific, Rockford, IL, USA) with 1x Protease & Phosphatase Inhibitor Cocktail (Thermo Scientific, Rockford, IL, USA) following the manufacturer’s protocol. Proteins were separated by 10% SDS-PAGE and transferred to a polyvinylidene difluoride (PVDF) membrane. Membranes were blocked using 5% nonfat dry milk in PBS containing 0.05% Tween 20, and then incubated with MITF (12590, Cell Signaling Technology, Danvers, MA, USA), DCT (SC-74439, Santa Cruz Biotechnology, Dallas, TX, USA), or GAPDH (SC-365062, Santa Cruz Biotechnology, Dallas, TX, USA) primary antibodies overnight at 4 °C. HRP-conjugated secondary antibody was used and detected using the Pierce ECL Western Blot Substrate. Blots were imaged with the ChemiDoc™ XRS+ Molecular Imager® (Bio-Rad Laboratories, Hercules, CA, USA).

### RT-qPCR

SK-MEL-30 or human primary melanocytes were treated with DMSO or A-485 (5 µM) for 24 hours. RNA was then extracted using the RNeasy (Qiagen, Germantown, MD, USA) kit and cDNA was synthesized using the SuperScript™ III Reverse Transcriptase kit (Invitrogen, ThermoFisher Scientific, Carlsbad, CA, USA) according to manufacturer instructions. qPCR was performed on the Applied Biosystems Step One Plus Real Time PCR System according to the manufacturer instructions using the iQ SYBR® Green Supermix (Bio-Rad Laboratories, Hercules, CA, USA) kit. Primer sequences are as follows: *18S* Forward: CTACCACATCCAAGGAAGCA, *18S* Reverse: TTTTTCGTCACTACCTCCCCG, *MITF* Forward: GGAAATCTTGGGCTTGATGGA, *MITF* Reverse: CCCGAGACAGGCAACGTATT, *DCT* Forward: TATTAGGACCAGGACGCCCC, *DCT* Reverse: TGGTACCGGTGCCAGGTAAC.

### Melanin Quantification

SK-MEL-30 or primary human melanocyte cells were treated with DMSO (control) or 5 µM A-485 for 6-days. Cells were lysed in 1 mL of 1 M NaOH and heated at 80°C for 2 hours. Samples were vortexed and subsequently centrifuged at 12,000xg for 10 minutes. The supernatant was collected, and melanin content was quantified by measuring absorbance at 405 nm and comparing measured values to a standard curve of melanin (M8631, Sigma-Aldrich, St. Louis, MO, USA) diluted in 1 M NaOH.

### Statistical analysis

Statistical significance was determined using a student’s *t*-test. A *p* value of <0.05 was considered statistically significant (^*^*p* < 0.05 and ^**^ *p* < 0.005). Standard deviation (SD) was utilized for error bars.

## Author Contribution Statement

Conceptualization: RA; Data Curation: NG, KL, ARW, AW; Formal Analysis: NAG, AWR; Funding Acquisition: RA; Investigation: NG, KL, AW, ARW, RA; Methodology: NG, ARW; Supervision: ARW, RA; Validation: NG, ARR, RA; Visualization: NG, ARW; Writing - Original Draft Preparation: NG, ARR, RA, MC; Writing - Review and Editing: ARW, RA, MC.

## Conflicts of Interest

RMA is a cofounder of Acylin Therapeutics and holds equity in the company. Acylin Therapeutics has an interest in developing epigenetic inhibitors, including A-485. RA and ARW have submitted intellectual property for utilizing A-485 to treat hyperpigmentation disorders through the Boston University Office of Technology Development.

## Acknowledgements

We thank Dr. Deborah Lang in the Department of Dermatology at Boston University for her expertise and guidance for melanin quantification.

